# Massively parallel reporter assays and mouse transgenic assays provide complementary information about neuronal enhancer activity

**DOI:** 10.1101/2024.04.22.590634

**Authors:** Michael Kosicki, Dianne Laboy Cintrón, Nicholas F. Page, Ilias Georgakopoulos-Soares, Jennifer A. Akiyama, Ingrid Plajzer-Frick, Catherine S. Novak, Momoe Kato, Riana D. Hunter, Kianna von Maydell, Sarah Barton, Patrick Godfrey, Erik Beckman, Stephan J. Sanders, Len A. Pennacchio, Nadav Ahituv

## Abstract

Genetic studies find hundreds of thousands of noncoding variants associated with psychiatric disorders. Massively parallel reporter assays (MPRAs) and *in vivo* transgenic mouse assays can be used to assay the impact of these variants. However, the relevance of MPRAs to *in vivo* function is unknown and transgenic assays suffer from low throughput. Here, we studied the utility of combining the two assays to study the impact of non-coding variants. We carried out an MPRA on over 50,000 sequences derived from enhancers validated in transgenic mouse assays and from multiple fetal neuronal ATAC-seq datasets. We also tested over 20,000 variants, including synthetic mutations in highly active neuronal enhancers and 177 common variants associated with psychiatric disorders. Variants with a high impact on MPRA activity were further tested in mice. We found a strong and specific correlation between MPRA and mouse neuronal enhancer activity including changes in neuronal enhancer activity in mouse embryos for variants with strong MPRA effects. Mouse assays also revealed pleiotropic variant effects that could not be observed in MPRA. Our work provides a large catalog of functional neuronal enhancers and variant effects and highlights the effectiveness of combining MPRAs and mouse transgenic assays.

## Introduction

Genome-wide association studies (GWAS) have identified hundreds of non-coding variants associated with psychiatric disorders, which exhibit complex genetic etiologies likely involving multiple loci^1–6^. The GWAS-discovered lead variants are not necessarily causative due to linkage disequilibrium (LD), which increases the number of potential variant candidates on average by ten-fold. In addition, ongoing whole genome sequencing studies of patients with psychiatric disorders identify 60-70 *de novo* non-coding variants per individual. These efforts highlight the challenge to distinguish causative variants from the hundreds of thousands of potential candidates identified through genetic studies.

Various genomic correlates of function can be used to reduce the number of potential candidates. Putative regulatory sequences can be identified in a tissue and even cell-type specific manner using such methods as DNase-seq and ATAC-Seq (for identification of open chromatin), or ChIP-seq^7–11^ (for identification of regions bound by transcription factors or having specific histone marks). Variants falling into regulatory regions with activity in relevant cell types are more likely to be causative. However, an overlap between a variant and regulatory region neither confirms variant functionality, nor provides a mechanism for how it impacts the phenotype. Functional assays that can test the effect of the variant on gene regulatory activity are needed to pinpoint the causative mutations.

Massively parallel reporter assays (MPRA) allow for the assessment of regulatory activity of tens of thousands to hundreds of thousands of candidate regulatory sequences and variants within them^12–14^. The majority of MPRAs are conducted *in vitro*, enabling the high throughput interrogation of candidate sequences and variants in a quantitative and reproducible manner. However, they are limited to testing the function of the assayed sequence only in the specific cell type and cannot assess how results relate to its function *in vivo*. As an alternative, *in vivo* activity of enhancers can be tested using a transgenic mouse assay (referred to as “transgenic assay” below) such as enSERT^15,16^. In this assay, a candidate regulatory sequence is coupled to a minimal promoter and reporter gene followed by its integration into a safe harbor locus in mouse zygotes and assayed for activity by imaging at a later embryonic time point. Transgenic assays can identify enhancer expression at an organismal level, providing rich, multi-tissue phenotype. Results of thousands of these assays are cataloged in the VISTA enhancer browser and serve as a gold standard for enhancer activity assessment^17^. However, these assays are more resource and labor intensive than MPRAs and therefore are typically conducted at a much lower throughput.

Combining the high throughput capabilities of MPRAs and rich phenotype of transgenic assays is an underexplored venue for regulatory element and variant characterization. Limited comparisons of these technologies have been performed^18–22^, but typically involved MPRAs conducted in cancer or immortalized cell lines with limited relevance to organismal biology, used short sequences (120 bp) or sampled too sparsely from *in vivo* validated sequences to enable a systematic comparison.

Here, we set out to robustly compare results between MPRA and transgenic assays by using psychiatric disorders-associated sequences and variants as a test case. We carried out an MPRA for over 50,000 sequences 270 bp in length, many of which were derived from brain enhancers in the VISTA enhancer browser^17^ and over 20,000 variants. We found thousands of functional regulatory sequences and hundreds of variants that alter regulatory activity compared to their reference allele. We observed an overall strong correlation between MPRA and transgenic assays. Variants with a high impact in MPRA also had a significant effect on neuronal enhancer activity in transgenic assays in mouse embryos. Combined, our work provides a large catalog of functional neuronal enhancers and their variants, and shows that MPRAs can be successfully combined with mouse transgenic enhancer assays.

## Results

### MPRA neuronal library composition and initial QC

We set out to investigate the correlation between high-throughput MPRAs and mouse enhancer transgenic assays. As neuronal enhancers are the most abundant category of enhancers in the VISTA Enhancer Browser^17^, which catalogs mouse transgenic assay results, and since our lab has established MPRA protocols in stem cell differentiated neurons^18,23,24^, we focused on neuronal-associated elements. We designed an MPRA library by tiling peaks from five single-cell and bulk neuronal ATAC-seq experiments^25–29^ and from conserved cores of 1,400 neuronal and non-neuronal enhancers from the VISTA Enhancer Browser^17^ with 270 bp tiles (Figure 1a,b; minimum 30 bp overlap; see Methods). To characterize how mutations affect the activity of these elements, we introduced two types of variants into the designed tiles. First, we included all lead single-nucleotide polymorphisms (SNPs) and SNPs in linkage disequilibrium (r^2^ > 0.8) from autism spectrum disorders (ASD), schizophrenia, bipolar disorders and depression GWAS that overlapped designed tiles^1,3–5^. Second, we introduced synthetic transversion variants into every fourth base pair of elements with high likelihood of MPRA activity (overlapping ATAC-seq peaks from multiple datasets, evolutionary conserved, active in transgenic assay, see Methods; Figure 1a,b)^30^. As negative controls, we used 500 di-nucleotide scrambled, non-conserved tiles from enhancers negative in mouse transgenic assays that did not have overlapping ENCODE candidate *cis*-regulatory elements^31^ or neuronal ATAC-seq signal^25–29^. In total, we designed 81,952 unique 270 bp sequences, including 24,942 variants.

**Figure 1.**
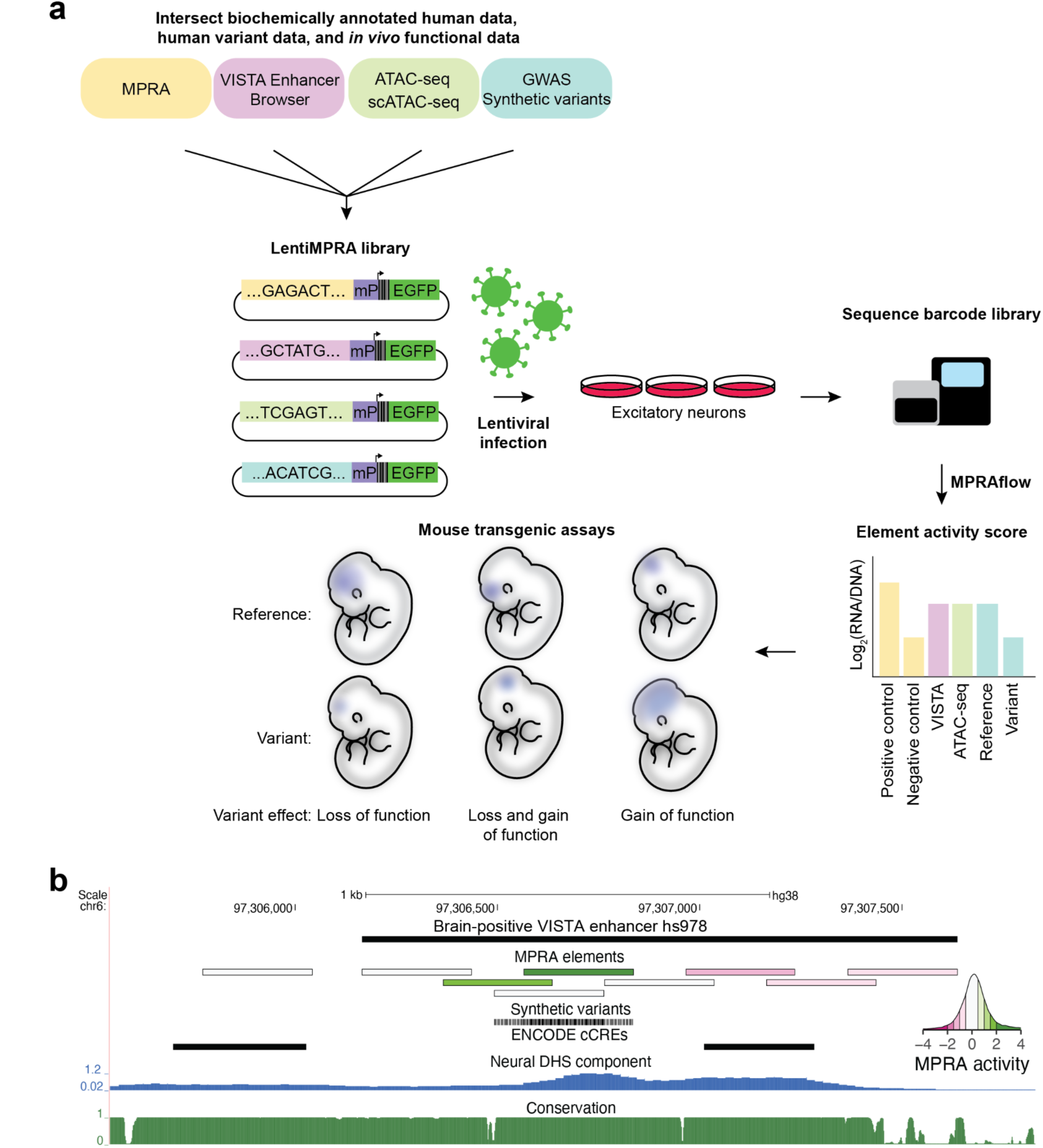
Functional validation of candidate *cis-*regulatory elements (cCREs) using lentiMPRA and mouse transgenic assays. **(a)** Schematic of experimental plan. A lentiMPRA library was designed through the intersection of scATAC-seq, ATAC-seq, VISTA Enhancer Browser^17^, conservation and neuronal MPRA data. The library also included GWAS lead SNPs and SNPs in LD with them and synthetic variants. Sequences were inserted into a reporter plasmid upstream of a minimal promoter (mP), barcode and EGFP. The library was infected into WTC11 induced excitatory neurons using lentivirus. The integrated DNA and transcribed RNA barcodes were sequenced to determine element activity scores. Mouse transgenic assays were conducted on selected sequences to characterize their activity *in vivo*. **(b)** UCSC Browser annotation, from top to bottom: (1) VISTA enhancer browser hs978 sequence (2) MPRA elements colored by MPRA activity with green showing high activity and pink low activity (see inset). (3) synthetic variants included in MPRA tested elements (4) ENCODE cCRE (candidate *cis*-regulatory elements)^31^ (5) Neural DNase I hypersensitivity signal component ^32^ (6) PhastCons conservation UCSC track for 30 mammals (27 primates).

Oligos were synthesized and cloned into a lentiMPRA vector and packaged into lentivirus following our previously published protocol^14^. They were then transduced into differentiated human excitatory neurons derived from an isogenic WTC11-Ngn2 iPSC line with an inducible Neurogenin-2 gene using an established induction protocol^14,33,34^. Only tiles with at least 15 barcodes detected in all three replicates were retained (median = 177 barcodes post-filtering) and tiles with mutations without a reference tile passing these criteria were discarded. Out of 81,952 elements, 76,415 passed QC (> 90%; see Methods), including 52,335 genomic elements, 23,482 single base pair mutation tiles and 476 scramble negative controls. MPRA activity was expressed as a z-score of log2(RNA counts/DNA counts) relative to scramble negative controls. We observed good correlation between replicates (Pearson correlation = 0.58-0.59, Supplementary Figure 1, N = 76,415). Based on nominal p-value < 0.05 from a t-test against mean of scramble negatives and an absolute activity at least least one standard deviation above scrambled controls, we designated 4,762 tiles to be activators and 2,957 tiles to be repressors (out of 52,335, 14.7%). Using similar criteria, we found 467 single base pair mutation tiles to have decreased activity compared to reference tile and 313 to have an increased one (out of 23,482, 3.3%).

### MPRA captures neuronal-specific activity

To validate the results of our MPRA, we annotated the activity of tiles overlapping a variety of genomic annotations. Specifically, we asked if ranks of the overlapping tiles were significantly different than scrambled negative controls (Figure 2a; Supplementary Table 1). On average, elements in our library were more active than scrambled negative controls (median activity = 0.19). Overlap with positive elements in previous neuronal MPRAs was associated with higher activity, with elements from Inoue 2019^18^ publication (double-Smad inhibition protocol) being more active than those from Uebbing 2021^35^ (stable neural stem cell line, median activity 0.31 vs 0.18). We also confirmed that tiles that were pre-selected for mutagenesis due to high expected activity were indeed highly active (“Mutation reference tiles”, activity = 0.33). At the positive extreme, tiles overlapping housekeeping promoters (defined as 2 kb centered around the 5’ end of Gencode protein-coding exon 1 of genes in Eisenberg and Levanon 2013^36^) were highly active (median activity 0.56), suggesting that they can be used as universal positive controls in other MPRAs. Ultraconserved elements^37^ had high activity as well (0.41), which is consistent with our previous observation that they are often active in developing brain in the transgenic assay^38^. Conversely, tiles overlapping coding exons, but not exons of long-non-coding RNAs, were overall repressive (median activity = -0.23). It is unlikely splicing sites present at exon-intron interface explain this observation, as in our MPRA the minimal promoter, and consequently the transcription start site, are downstream of the tested element. In addition to rank-based analysis, we also checked if the proportion of activators to repressor tiles overlapping different genomic annotations was significantly different from non-overlapping ones. The results obtained using this method were similar (Supplementary Figure 2a; Supplementary Table 1).

**Figure 2.**
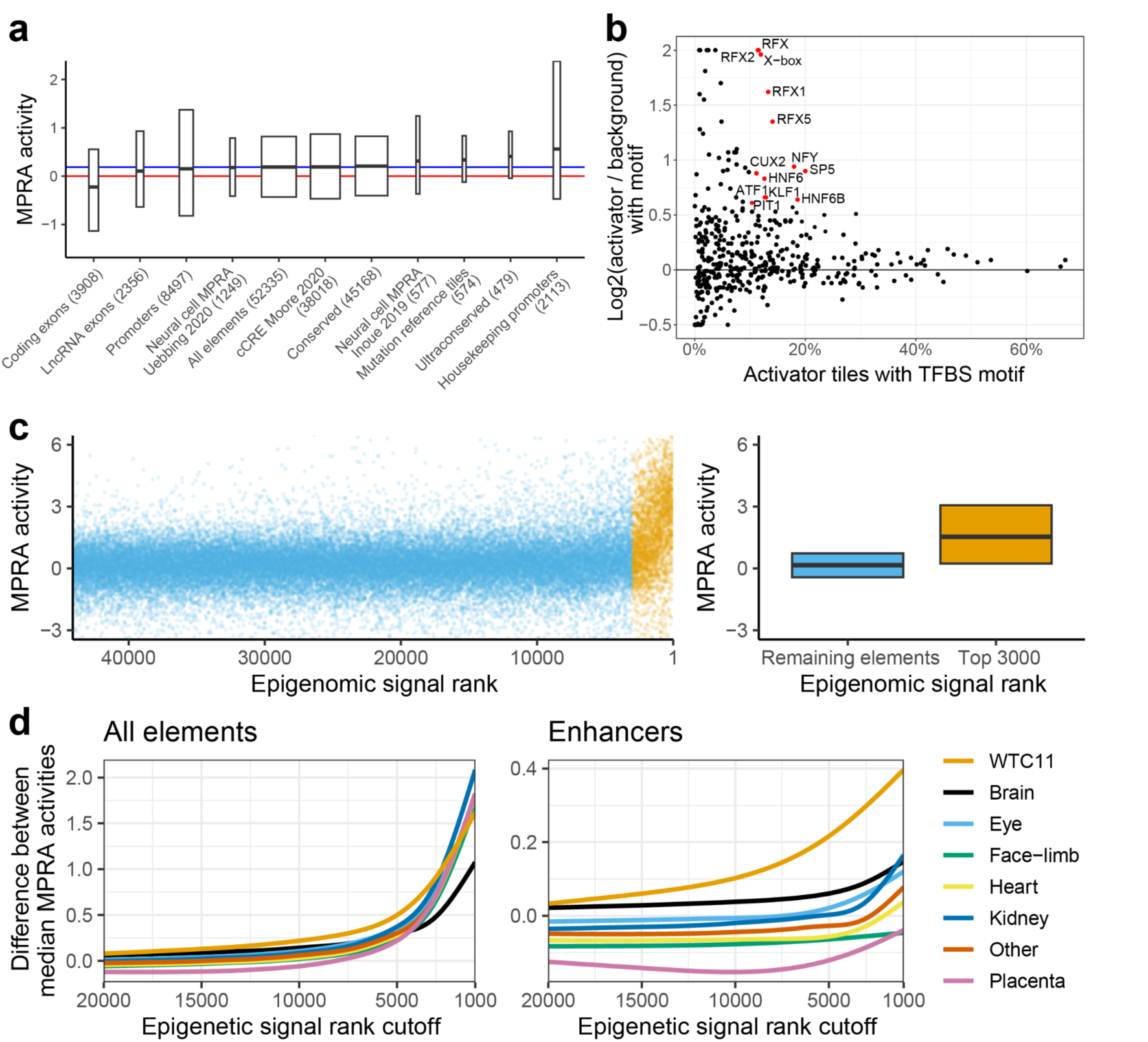
Neuronal WTC11 MPRA results validation. **(a)** MPRA activity of tiles overlapping different categories. Red line = activity of scrambled negative controls (zero, by definition). Blue line = median activity of all reference elements (0.19). Hinges of boxplot span interquartile range, line in the middle is median, width is proportional to the number of overlapping tiles. All categories have significantly different activity than scrambled negatives at FDR-adjusted p-value < 0.05 (Mann-Whitney U test). Promoters = 5’ end of exon 1 of protein-coding genes +/-1 kb. **(b)** TFBS enrichment in enhancer, activator tiles compared to enhancer elements with scramble negative levels of activity. Log2-fold change was curbed at -0.5 and 2. Only TFBSs present in more than 10% target, with 50% increase in presence from background to target set (corresponding to log2(1.5) = 0.58 cutoff) and FDR < 1% are labeled and colored red. **(c)** Methodology for comparison of epigenomic annotation. Left: tiles were ranked by epigenomic signal and split at various rank cutoffs into two groups. Right: Median MPRA activity of the two groups was compared. Same boxplot display conventions as in (a). **(d)** Difference in median MPRA activity at different epigenetic rank cutoffs for eight tissue groups. Left: all elements (N = 44,109; lower than 52,335 total due to exclusion of elements that failed to lift-over between human and mouse genomic assemblies), right: enhancers (N = 37,813; enhancers defined as not overlapping “coding promoter” category in (a); see Methods).

We then set out to analyze the transcription factor binding sites (TFBS) that could be involved with enhancer neuronal activity. Using HOMER^39^, we analyzed human and mouse TFBS that are enriched in activator tiles that do not overlap promoters (activity > 1, p-value < 0.05, N = 3,054) with tiles with background level activity, controlling for GC-content (-0.4 < activity < 0.4, N = 15,503; Figure 2b; Supplementary Table 2). We considered a TFBS to be enriched if it was present in at least 10% activator tiles, increased by at least 50% compared to background set (corresponding to log2(1.5) = 0.58 cutoff) and was significantly enriched by HOMER’s hypergeometric test at FDR-adjusted p-value < 0.01. We found a total of 13 motifs to be enriched in activator tiles, including neuron-associated motifs SP5, KLF1, CUX2 and five motifs from the RFX family^40,41^, as well as a NFY-binding promoter motif CCAAT, two liver, pancreas and nervous system expressed HNF6/ONECUT1 motifs^42^, pituitary-development associated PIT1/POU1F1 (POU family) and growth/survival TF ATF1. Analogous analysis using GC-matched genomic background found the same motifs (except ATF1 and KLF1) and a score of additional, neuronal-associated ones, mostly from SOX, LHX, DLX and E-box families (NEUROD1, MYOD, ATOH1; Supplementary Figure 2b,c)^41,43^. Repressor tiles that do no overlap promoters (activity < -1, p-value < 0.05, N = 1,894) were enriched for similar motifs as activators, when compared to genomic background. In particular, we observed SOX, LHX and DLX families, with only repressor-specific hits being SOX3 and BORIS/CTCFL (Supplementary Figure 2b,c). No repressor-enriched motifs were found when comparing to tiles with background-level activity.

We observed that both activator and repressor tiles had higher median levels of GC-content than the rest of the library, with repressors having higher levels than activators (repressors 64%, activators 50%, remaining elements 44%; Supplementary Figure 2d). Such GC-skew should not affect MPRA readout on a technical level, as the activity of the tested element is read through sequencing of a barcode, not the element itself (unlike e.g. in STARR-seq). We conclude that highly GC-rich sequences often function as transcriptional repressor elements in this MPRA.

We next set out to assess how well various biochemical marks correlated with neuronal MPRA activity. We compared our MPRA results to epigenomic signal from 12 embryonic, fetal and WTC11 datasets, encompassing 740 Dnase hypersensitive sites (DHS), ATAC and single-cell ATAC samples, covering diverse tissues and cell types (Supplementary Table 3). To account for a large diversity of experimental and computational protocols, we integrated raw genomic signal (bigWig tracks) over MPRA tiles and ranked the tiles based on the signal for each dataset. We then computed the difference between median MPRA activity of top ranked elements and the remaining ones for a range of epigenomic rank cutoffs (Figure 2b). As expected, the more stringent the rank cutoff, the larger the difference between activity of top ranked elements versus the rest. However, due to enrichment of promoter-overlapping elements in top ranks, the differences between individual datasets was negligible (Figure 2c, left). After removing tiles overlapping protein-coding promoters, we observed a clear separation of brain and differentiated WTC11 cells samples from non-neuronal samples (Figure 2d, left). Closer inspection revealed that some non-neuronal ENCODE DHS samples (adrenal, eye and kidney) are still enriched, especially at stringent cutoffs, possibly reflecting a combination of high activity tissue-invariant elements (“housekeeping enhancers”) and higher signal-to-noise ratio of DHS data at high signal intensities. This was attenuated at less stringent signal cutoffs, with only four non-neuronal samples (eye and adrenal) remaining in top 50 at signal rank cutoff 5000 (Supplementary Table 3). Encouragingly, we observed a clear separation over ATAC-seq time course of WTC11 cell neuronal differentiation, with undifferentiated cells ranking at position 599, day 3 differentiated cells at position 62 and day 14 at position 7^44^. We note that our MPRA design sampled elements with open chromatin signal in neuronal tissues more deeply than in other tissues, which may have contributed to the observed enrichments. In summary, our results show that our MPRA captures neuronal-associated regulatory activity.

### Neuronal MPRA activity correlates with mouse neuronal enhancer expression

The average sequence length tested in transgenic mouse assays is around 1 kb, around four times the size of tiles in our MPRA (270 bp). To compare these two assays, we matched transgenic assay elements (“VISTA elements”) with overlapping MPRA tiles of highest activity (Figure 3a). We then built a general linear model (GLM) with a binomial link to predict binary, tissue-specific transgenic assay results (e.g. brain activity, yes or no) from MPRA activity. Our coverage of VISTA elements in MPRA was biased towards conserved sequences of neural-positive elements, while avoiding these in negative elements (to create negative control tiles; Figure 3a). To account for that, we included a fraction of conserved sequences covered by tiles as a covariate in the model (Figure 3b,c). An alternative solution, removing poorly covered VISTA elements yielded similar results (Supplementary Figure 3).

We found that all neural annotations (except dorsal root ganglion) were significantly correlated with MPRA activity, while craniofacial and heart terms were significantly anti-correlated (Figure 3c). The steepest regression slope on the positive side was for a combined ‘neural’ term (brain, neural tube, cranial nerves including trigeminal nerve and dorsal root ganglion), followed by ‘brain’. The model predicted that to achieve a validation rate of 70%, which is typically the desired rate of transgenic assays, the tested element needs to overlap a tile with MPRA activity of at least 1.32. We conclude that neural MPRA in differentiated human excitatory neurons and neural activity in transgenic assay strongly correlate in a tissue-specific manner.

**Figure 3.**
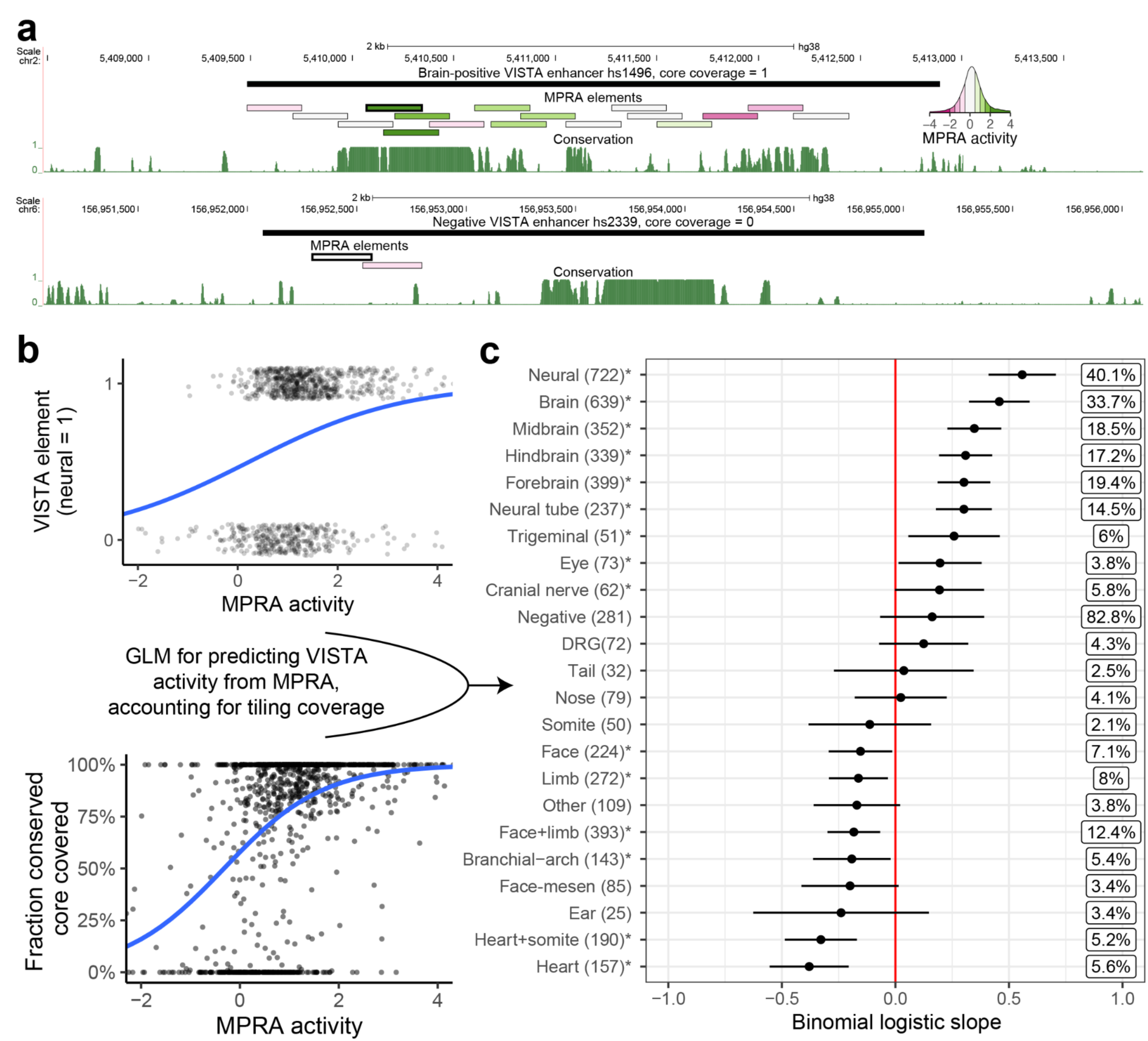
Predicting transgenic assay activity using a MPRA-based, coverage-corrected model. **(a)** Examples of VISTA elements with complete (top) and zero (bottom) coverage of their conserved cores using MPRA tiles. Conservation is PhastCons UCSC tracks for 30 mammals (27 primates) dataset. MPRA tile with highest activity (used for modeling) has a thicker border. MPRA elements colored by MPRA activity, see inset. **(b)** Visualization of input variables for the GLM. Top: transgenic assay (VISTA) elements are binarized according to chosen tissue activity (here: neural). The blue line is the binomial-link GLM regression on this variable. Bottom: relationship between fraction of conserved core covered and MPRA activity is modeled as a covariate. The blue line is the binomial-link GLM regression on this variable. **(c)** Results of the GLM predicting binomial transgenic assay activity from MPRA activity and fraction of conserved core covered. Asterisks indicate nominal p-value < 0.05. Boxed percentages to the right are Nagelkerke R^2^ measures. DRG = dorsal root ganglia. Face-mesen = facial mesenchyme. Cranial nerves category does not include the trigeminal nerve, as per VISTA Browser.

### Minimal MPRA effect of psychiatric disorder associated GWAS variants

We next analyzed the 177 psychiatric disorder associated GWAS variants tested in our MPRA. Using nominal significance criteria (see Methods), we found that only 3 out of the 177 variants had a significant effect on MPRA activity (Supplementary Table 4). Each variant was associated with an independent GWAS signal (different lead SNPs, two associated with bipolar disorder, one with major depressive disorder) and had a moderate, gain-of-activity impact on expression (1.1-1.5 units). None of the variants remained significant after multiple testing adjustment, all were in tiles with either repressive or no activity (-1.79 to 0.13) and tended to be outside or at the edge of the peak of the DHS signal and conservation (Supplementary Figure 4, Supplementary Table 4). Therefore, we decided not to further investigate them using the transgenic assay.

### Variants altering MPRA activity affect neuronal mouse enhancer activity

To select variants for transgenic assay follow-up, we analyzed the synthetic, single nucleotide variants. Out of 23,266 tiles with non-GWAS variants, 777 had a nominally significant effect on regulatory activity (p-value < 0.05, absolute log2 fold-change > 1). We prioritized variants with links to important neuronal genes predicted using ABC^45^ (e.g. *QKI*, *PRKN*, *COA7*, *SETBP1* and *MEF2C*) and prior evidence of neuronal activity in transgenic assays. Ultimately, we selected seven loss-of-function variants with different MPRA effect sizes (1-4.5 units; all significant, except one, see Table 1) and tested their impact on *in vivo* neural enhancer expression using a transgenic assay (Figure 4a).

**Table 1.** MPRA variants tested in transgenic mouse assay. TFBS predictions from motifbreakR. Motifs consistent with flanking variant effects in bold (see Figure 4c). Neural target genes found using activity-by-contact (ABC) on data from WTC11 excitatory neurons (cell line used in this study) or prenatal week 18 prefrontal cortex neurons (Methods). Asterisk - variant effect not significant.

| Variant name | Reference MPRA activity | Variant MPRA effect | TFBS affected | VISTA element | Structures affected in transgenic assay | Neural target genes | Relevant target gene Phenotypic associations |
| --- | --- | --- | --- | --- | --- | --- | --- |
| chr6-97306759-A-T | 4.1 | -4.5 | **CDX1 (gain), POU4F1** | hs978.1 | partial forebrain, hindbrain and neural tube loss | GPR63 | Branchiooculofacial Syndrome and Spina Bifida Occulta |
| chr6-162856979-C-G | 3.8 | -4 | **RORB (gain), CTCF** | hs2793.1 | partial forebrain loss | QKI, PACRG and PRKN | Associated with Parkinson's Disease (PACRG, PRKN) and schizophrenia (QKI) |
| chr1-52663061-C-G | 6.2 | -4.5 | **SP2** | hs2790.1 | midbrain gain, partial forebrain loss | COA7, TUT4 and others | Spinocerebellar Ataxia (COA7) Perlman Syndrome (TUT4) |
| chr18-44826694-C-G | 0.6 | -2.6 | **MAFB** | hs2791.1 | partial cranial nerve loss, midbrain, hindbrain and neural tube gain | SETBP1 | Autism |
| chr4-104425251-C-G | 1.2 | -1* | **SATB1, RELA** | hs2792.1 | gain of unidentified forebrain or craniofacial structure | CXXC4 (IDAX) | Overexpression in frog embryos causes ventralization |
| chr5-88396638-C-G | -0.3 | -2.5 | **ASCL1 (gain), NR2F1** | hs268.1 | no effect | MEF2C | Associated with cognitive disability, epilepsy and cerebral malformations |
| chr5-88397333-A-T | 0.6 | -1.5 | [none] | hs268.2 | no effect |  |  |

We found that 5/7 variants affected mouse enhancer expression in a reproducible manner, with 4 causing a loss of activity in different parts of the brain, neural tube or cranial nerves (Figure 4b). In two cases, this was accompanied by a gain of expression in another brain-associated structure. One variant caused a gain of activity without any loss of function, in a suspected forebrain or craniofacial structure (hs2792.1). That variant was the only one not to pass our significance criteria in MPRA. We note that the two variants with no apparent impact on transgenic enhancer activity had a very high basal activity of the reference element in the transgenic assay (hs268), which may have masked expression differences due to the variant. These results demonstrate the utility of combining the two experimental systems, with a good correlation between MPRA and mouse transgenic assay and rich additional information provided by the latter.

To further interpret the results of our transgenic assays, we used motifbreakR^46^ to carry out TFBS predictions for all seven variants tested using transgenic assay. We found plausible hits for all variants except hs268.2, which did not have an effect in the transgenic assay (Figure 4c,d; Supplementary Figure 6). In four cases, more than one plausible TFBS was found. We leveraged the fact our MPRA design also contained variants in close proximity to the ones we selected for transgenic assay follow up to further validate the TFBS predictions. For example, the variant tested in hs2792.1 element was predicted to affect both SATB1 and RELA binding. The flanking variant MPRA effects were consistent with SATB1 binding i.e. the flanking variant that reduced MPRA activity was at a position crucial to the TF motif, while the flanking variant with no impact on MPRA activity was at a position of low information content Figure 4c, hs2792.1). Conversely, RELA binding was not consistent with two of the flanking variants. The first of these variants had low predicted relevance for RELA binding (based on RELA motif), but exerted strong loss-of-function effect on MPRA activity (last T>A variant in Figure 4c). The second variant had no impact on MPRA activity, but was at a position crucial for RELA binding. While the first discrepancy could potentially be explained by binding of another, undetected TF, the other one is inconsistent with proposed RELA binding. We applied similar logic to remaining predictions to select the more plausible of the initial TFBS matches. Deploying this approach in a systematic manner could help interpret future variant MPRAs.

**Figure 4.**
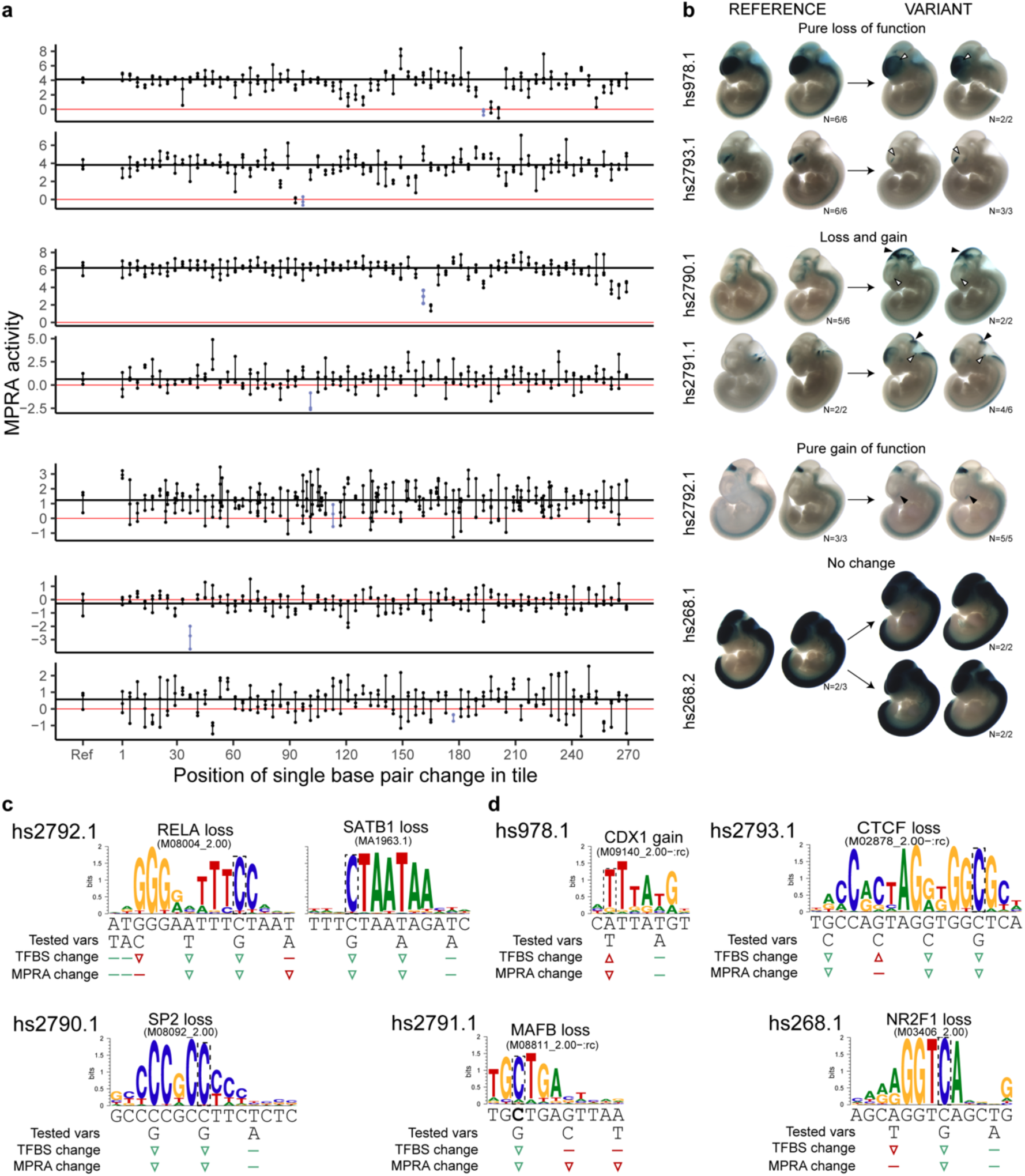
Synthetic MPRA variants lead to *in vivo* change of function in transgenic assay. (a) Every fourth nucleotide of seven MPRA tiles was mutagenized individually, for a total of 67 mutagenized constructs per reference tile (for tile related to hs2792.1 more variants were tested). Dots connected by a vertical line = three biological replicates. Red horizontal line = zero, mean activity of scramble negative controls. Black horizontal line = mean activity of the reference construct. **(b)** Constructs encompassing the MPRA tiles, with or without the variants indicated in blue in panel A, were tested for activity using transgenic assay in developing mouse embryos at embryonic day e11.5. All variants except hs268.1 and hs268.2 led to change of function in one or more neural tissues - brain, neural tube or cranial nerves. Shown are embryos that were genotyped as “tandem”, i.e. positive for insertion at the safe harbor locus and presence of the plasmid backbone indicating multi-copy insertion with strong, reproducible pattern. White arrowhead indicates loss of function, black indicates gain of function. See Supplementary Figure 5 for all embryo images, which provide additional support of the changes observed when comparing tandem embryos. **(c)** Prediction of TFBS likely affected by the variants for hs2792.1. Left: TFBS inconsistent with flanking effects (RELA), right: TFBS consistent with flanking variant effects (SATB1). TFBS and MPRA change symbols are colored green if matching and red if not. Arrowheads indicate an increase or decrease, flat line indicates no effect **(d)** Prediction of TFBS likely affected by the variants and partially or fully consistent with flanking variant effects.

## Discussion

We performed an MPRA in neurons with elements derived from VISTA enhancers and neuronal fetal ATAC-seq peaks finding a good correlation to neuronal expression in mouse transgenic assays. In terms of variants, we did not see a strong effect on MPRA activity for our selected psychiatric disorder associated GWAS variants, but observed effects on MPRA activity for 777 out of 23,266 synthetic variant tiles. Most synthetic variants nominated by MPRA as having a large impact also affected the transgenic assay activity in the expected manner and revealed additional ectopic effects. Overall, we demonstrated that combining MPRA and transgenic assays can be highly advantageous.

The observed complementary of the two assays is encouraging. MPRA allows the testing of a large number of sequences and provides a quantitative readout, while transgenic assays can reveal the organismal spatially-resolved consequences of regulatory sequences and variants. Both approaches have improved significantly over the past decade, coming closer to bridging the gap between them. MPRAs have been increasing in throughput, length of tested elements, range of cell types amenable to this type of assay (due to lentivirus and AAV delivery) and have also been carried out *in vivo* in select tissues in a postnatal manner^14,21,22,48–50^. Transgenic assays improved in throughput and reproducibility due to the development of Cas9-guided safe harbor integration method enSERT^16^. Their remaining shortcomings can be overcome by combining the techniques. MPRAs conducted *in vitro* are limited to the cell types in which they are assayed, can be limited by the availability of differentiation protocols and labor intensity of differentiating millions of cells with various identities and cannot assess the spatial and temporal organismal activity of a regulatory element. enSERT is conducted in mice, which cannot capture all aspects of human biology, is costly and not high-throughput. It also has only recently been applied in a quantitative manner^51^. While both methods are likely to improve and eventually merge, our work highlights the utility of combining currently available approaches, with MPRA as a high-throughput filter for the multi-tissue transgenic assay.

We did not observe a significant effect on MPRA activity for the 177 tested psychiatric disorder associated GWAS variants after applying multiple testing correction. This is in line with another MPRA carried out by our lab that found only 164 psychiatric disorder and eQTL variants out of 17,069 tested (< 1%) to have an effect on MPRA activity^50^. This could be due to a variety of reasons: 1) The small number of variants tested in our study; 2) Generally low expected effect size of common variation associated with complex traits such as psychiatric disorders. Machine learning models of MPRA data^50^ suggest that rare variants have a higher effect on MPRA activity compared to common variants; 3) Some variants may affect non-transcriptional phenotypes, like chromatin tethering; 4) Some variants may have an effect in another cell type or at a different differentiation time point.

Synthetic variants comprised the majority of variants tested in this MPRA, which has some advantages over testing common variants. First, the effect sizes of these variants are not constrained by negative selection, unlike common variation in human populations. This makes synthetic MPRA a better substrate for computational modeling, which should be able to learn a wide range of potential effects. Second, our experiment allowed us to find functional variants in elements likely to control expression of neuronal genes, some of which are linked to neurodevelopmental disorders. These results place a strong prior on interpretation of yet undiscovered, large effect de novo variants in these regions and can help better understand the regulatory biology of neuronal development.

In summary, we compiled a catalog of transcriptional activity in neuronal cells of over 50,000 elements derived from open chromatin fetal datasets and enhancers validated in transgenic assays. We also assessed the impact of over 20,000 synthetic and 177 GWAS variants, and demonstrated the usefulness of using MPRA as a variant filter for transgenic mouse assays. We anticipate this work will contribute to computational modeling of gene regulation and studies focused on neural development and psychiatric disorders.

## Methods

### MPRA design

We used following datasets for our library design - VISTA enhancers, MPRA tiles from Inoue 2019^18^ (activity > 1.1 at both 48h and 72h timepoints) and Uebbing 2021^35^ (q<0.05 for both replicates, following the publication) and single-cell or bulk ATAC or ATAC-seq peaks called by Ziffra 2020^25^ (26,000 peaks designated enhancers by activity-by-contact), Domcke 2020^26^ (33,000 cerebrum peaks with mean expression > 0.1), Preissl 2018^27^ (top 15,000 peaks from each of eEX1, eIN1, eIN2 and RG1-4 clusters), Gorkin 2020^28^ (top 15,000 from forebrain, midbrain, hindbrain and neural tube e11.5 samples), Inoue 2019^18^ (top 15,000 peaks from 72h timepoint) and Song 2019^29^ (WTC11 neurons; top 15,000 peaks; Supplementary Table 5). These elements were either extended to 270 bp (if shorter) or tiled in intervals of 270 bp with a minimum 30 bp overlap. We also designed tiles directly upstream of first exons of coding genes in Gencode v34 with neural ATAC signal (one tile per promoter), facing in the direction of transcription and avoiding overlap with FANTOM5 CAGE TSS peaks^52^. Tiles centered on rDHS elements with ‘Neural’, ‘Organ devel. / renal’ and ‘Primitive / embryonic’ annotation were added, if overlapping previously chosen elements^32^. Genomic negative control tiles (N = 500) were selected from sections of negative VISTA elements that were not conserved, not active in previous neural MPRAs and did not overlap any cCREs^31^ or any of the peaks in ATAC datasets mentioned above. Finally, we used a weighted combination of evolutionary conservation (UCSC phastConsElements30way^53^), lack of overlap with coding exons (Gencode^54^) and promoters regions (Gencode and CAGE^52^), neural VISTA activity, presence of a peak in multiple ATAC datasets, activity in previous neural MPRAs^18,35^ and overlap with LD blocks from psychiatric disorder GWAS to select 56,387 hg38 genomic reference tiles. We also used di-nucleotide scrambled 500 genomic negative control tiles to form scramble negative controls. All resources that were not originally available in hg38 (including mouse VISTA enhancers), were lifted over using Kent tools and relevant UCSC chain files^55^. Design was conducted in R 4.3.2 with tidyverse 2.0.0 package^56,57^. Additional information about the used data and software can be found in Supplementary Table 5.

We introduced mutations into 595 reference tiles, resulting in 123 tiles with multiple SNVs (derived from random mutagenesis of ultraconserved VISTA elements^58^) and 24,942 with individual SNVs. To select GWAS SNPs for testing, we started with a set of 465 common lead SNPs from psychiatric disorder GWAS^1,3–5^ and extracted 15,133 SNPs in linkage disequilibrium (LD) with these using SNiPA (r^2^ > 0.8, 1000 genomes set, v3). We selected 186 for testing based on overlap with enhancers with high likelihood of activity (overlapping ATAC-seq peaks from multiple datasets, evolutionary conserved, active in transgenic assay or highly active in previous neuronal MPRAs). To investigate vulnerabilities of regulatory elements associated with GWAS signals, we conducted systematic mutagenesis of every fourth nucleotide in 85 tiles within 0.8 LD regions for additional 5,621 SNVs using a GC-preserving transversion scheme (G=C, A=T). Finally, we conducted similar systematic mutagenesis of 254 tiles with high likelihood of activity for an additional 17,272 GC-preserving transversion SNVs. Numbers of SNVs listed above are mutually exclusive, but some SNVs belonged to more than one category. For example, a total of 5,892 SNVs were in 0.8 LD regions, including synthetic, lead and LD SNPs. Note that about 10% of all designed elements were not successfully tested - see Results section for numbers after QC.

### LentiMPRA cloning and infection

The lentiMPRA library was constructed as previously described^14^. A synthesized TWIST oligo pool with 300 bp long elements (270 bp insert + 2*15 bp PCR handles) was amplified by 12-cycle PCR using NEBNext High-Fidelity 2x PCR Master Mix (New England BioLabs, M0541L), the forward primer 5BC-AG-f01 and reverse primer 5BC-AG-r01 were used to add the minimal promoter, spacer and vector overhang sequence. The amplified fragment was purified using 1x of the HighPrep PCR Clean-up System (Magbio, AC-60500). The purified fragment underwent a second round of 12-cycle PCR using NEBNext High-Fidelity 2x PCR Master Mix (New England BioLabs, M0541L), the forward primer 5BC-AG-f02 and the reverse primer 5BC-AG-r02. This step added a 15 bp random barcode downstream from the minimal promoter. The amplified fragment was purified using Nucleospin Gel and PCR-Clean-Up (Macherey-Nagel, 740609.50) and 1.2x HighPrep PCR Clean-up System (Magbio, AC-60500). The oligo library was cloned into the double digested AgeI/SbfI pLS-SceI vector (Addgene,137725) using the NEBuilder HiFi DNA Assembly Master Mix (New England BioLabs, E2621L). The plasmid lentiMPRA library was electroporated into MegaX DH10B T1R Electrocomp Cells (Invitrogen, C640003) using the Gemini X2 (2.0 kV, 200 Ω, 25 µF). The electroporated cells were then plated on eleven 15 cm 100 mg/mL ampicillin LB agar plates (Teknova, L5004) and grown overnight at 37 °C. Approximately 8 million colonies were pooled and midi-prepped (Qiagen, 12145) to obtain on average 100 barcodes per oligo. To associate barcodes with each oligo in the library, the Illumina flow cell adapters were added through a 15-cycle PCR using NEBNext High-Fidelity 2x PCR Master Mix (New England BioLabs, M0541L), the forward primer pLSmP-ass-i741 and reverse primer pLSmP-ass-gfp. The amplified fragment was purified using Nucleospin Gel and PCR-Clean-Up (Macherey-Nagel, 740609.50) and 1.8x HighPrep PCR Clean-up System (Magbio, AC-60500). The amplified fragments were sequenced on a NextSeq 300 using a NextSeq 150PE kit with custom primers (R1: pLSmP-ass-seq-R1, R2: pLSmP-ass-seq-ind1, R3: pLSmP-ass-seq-R2).

Lentivirus production was conducted on twenty-nine 10 cm dishes of LentiX 293T cell line (TakaraBio, 632180) with Lenti-Pac HIV expression packaging kit (GeneCopoeia, LT002) following the manufacturer’s protocol. Lentivirus was filtered through a .45 µm PES filter system (Thermo Fisher Scientific,165-0045) and concentrated with Lenti-X Concentrator (TakaraBio, 631232). Titration of the lentiMPRA library was conducted on differentiated human excitatory neurons. Cells were seeded at 4.5 x 10^4^ cells per well in a 12-well plate on day 0 and incubated for 7 days. Serial volumes of the lentivirus (0, 1, 2, 4, 8, 16, 32, 64,128 µL) were added along with 6 µL ViroMag R/L (OZ Biosciences, RL41000) per well. After lentivirus addition cells were incubated for 30 minutes on the magnet at 37 °C. The magnet was removed and cells were incubated at 37 °C for 7 days, the media was replaced after 24 hours of lentivirus addition. The cells were washed with DPBS (Sigma-Aldrich, D8537) and DNA was extracted with the AllPrep DNA/RNA Mini Kit (Qiagen, 80204) following the manufacturer’s protocol for DNA extraction.

The multiplicity of infection (MOI) was determined as the relative amount of viral DNA over that of genomic DNA by qPCR using SsoFast EvaGreen Supermix (Bio-Rad, 1725205). The lentivirus infection, DNA/RNA extraction and DNA/RNA barcode sequencing were conducted as previously described^14^. Each replicate required approximately 25 million cells. Therefore, cells were seeded at day 0 of differentiation in four 10 cm plates with 5 million cells each. On day 7, the cells were infected with the lentivirus library and ViroMag R/L (OZ Biosciences, RL41000) following the manufacturer’s protocol. All three replicates were infected with the same lentivirus batch at an MOI of 80. Media was replaced 24 hours after lentivirus addition and the cells were incubated for 7 days. DNA and RNA were extracted from the three replicates using the AllPrep DNA/RNA Mini Kit (Qiagen, 80204) following the manufacturer’s protocol. The RNA was treated with the TURBO DNA-free Kit (Life Technologies, AM1907) following the manufacturer’s protocol for rigorous DNase treatment. Reverse transcription was conducted with SuperScript II Reverse Transcriptase (Life Technologies, 18064-071) using a barcode-specific primer (P7-pLSmP-ass16UMI-gfp) which contains a 16 bp UMI. After DNAse treatment and reverse transcription the resulting cDNA and extracted DNA underwent the same steps to prepare the library for sequencing. To add a sample index and UMI, DNA and cDNA from the three replicates were kept separate and underwent a 3-cycle PCR using NEBNext Ultra II Q5 Master Mix (New England Biolabs, M0544L), forward primer P7-pLSmp-ass16UMI-gfp and reverse primer P5-pLSmP-5bc-i#. Another round of PCR was conducted to prepare the library for sequencing using NEBNext Ultra II Q5 Master Mix (New England Biolabs, M0544L), forward primer P5 and reverse primer P7. The fragments were purified using 1.2x of the HighPrep PCR Clean-up System (Magbio, AC-60500). The final libraries were sequenced on four runs of Illumina NextSeq high-output using the custom primers (R1: pLSmP-ass-seq-ind1, R2: pLSmP-UMI-seq, R3: pLSmP-bc-seq, R4: pLSmP-5bc-seq-R2).

### Cell culture and neuronal differentiation

Differentiated human excitatory neurons were derived from hiPSCs in the WTC11 background where a doxycycline-inducible neurogenin 2 transgene was integrated into the AAVS1 locus^33^. In the undifferentiated stage, cells were maintained in mTeSR 1 (STEMCELL Technologies, 85850) and the medium was changed daily. Once confluent, cells were washed with 1x DPBS (Sigma-Aldrich, D853), dissociated with accutase (STEMCELL Technologies, 07920) and plated a at 1:6 ratio in matrigel (Corning, 354277) coated plates. Media was supplemented with ROCK inhibitor Y-27632 (STEM CELL, 72304) at 10 µM on the day of passage. To initiate differentiation, cells were washed with 1x DPBS, dissociated with accutase and plated in matrigel-coated plates. For three days cells were cultured in KnockOut DMEM/F-12 (Life Technologies, 12660-012) medium supplemented with 2 µg/mL doxycycline (Sigma-Aldrich, D9891), 1X N-2 Supplement (Life Technologies, 17502-048), 1X NEAA (Life Technologies, 11140-050), 10 ng/mL BDNF (PeproTech, 450-02), 10 ng/mL NT-3 (PeproTech, 450-03) and 1µg/mL lamininin (Life Technologies, 23017-015). The pre-differentiation medium was changed daily for three days and on the first day medium was supplemented with 10 µM ROCK inhibitor Y-27632. To induce neuronal maturation, cells were lifted and plated in Poly-L-Ornithine (Sigma-Aldrich, P3655) coated plates. Cells were cultured in maturation media containing Neurobasal A (Life Technologies, 12349-015) and DMEM/F12, HEPES (Life Technologies, 11330-032) with 2 µg/mL doxycycline supplemented with 1X N-2 Supplement, 0.5X B-27 Supplement, 1X NEAA, 0.5X GlutaMax (Life Technologies, 35050-061), 10 ng/mL BDNF, 10 ng/mL NT-3 and 1 µg/mL lamininin. A half-media change was conducted on day 7 and day 14 of differentiation using the maturation medium minus doxycycline.

### LentiMPRA analysis

Processing of barcode association and final MPRA libraries was done using a standardized MPRAflow pipeline^14,34,59^, without a MAPQ filter to avoid artificial dropout due to multi-mapping of elements with single base pair mutations. All subsequent analyses were conducted in R 4.3.2 with tidyverse 2.0.0 package^57^. Visualizations were done using ggrastr 1.0.2 (https://github.com/VPetukhov/ggrastr), ggplot2 3.5.0^60^ and ggrepel 0.9.5^61^. General linear models were constructed using rms 6.8-0 (https://hbiostat.org/r/rms/). Motifs affected by variants tested in the transgenic assay were detected using motifbreakR 2.16.0^46^ with filterp=T, threshold=1e-4 and pwmList from Viestra 2020^62^. Only tiles with at least 15 barcodes detected in all three replicates were retained and mutation tiles without a reference passing these criteria were discarded as well. MPRA activity was expressed as a z-score of log2(RNA counts/DNA counts) relative to scramble negative controls.

### Correlation of MPRA activity and epigenomic signal

Epigenomic signal in the form of bigWig files was retrieved from ENCODE^31^ and 12 other sources (Supplementary Table 5) and integrated over tile intervals using *bedtools bigWigAverageOverBed* command^63^. For each sample, signal was sorted and ranked with random tie-breaking. For a range of rank cutoffs starting with 1000, the tiles were split into those above and below the cutoff and median MPRA activity was computed for both groups. The median activity of bottom signal group (e.g. from rank 1001 to lowest rank) was then subtracted from median activity of the top signal group (e.g. ranks 1-1000). For enhancer analysis, 8495 tiles overlapping promoters defined as 2 kb centered on the 5’ end of exon 1 of protein-coding genes in Gencode V34^54^, were removed before computing the ranks and median activity difference.

### TFBS enrichment analysis

All analysis was done using HOMER 4.11^39^ using activator or repressor tiles as target (as defined in the main text) and either HOMER-selected, GC-matched background genomic elements of the same size, or library elements with scramble negative levels of activity (-0.4 < activity < 0.4). Only tiles not overlapping promoters, as defined in the previous section, were used. Default set of 239 unique TF motifs was used. Command of the form *findMotifsGenome.pl target.bed hg38 target_folder -bg background.bed -size 270 -nomotif* was run for each analysis, except *-bg* term was dropped for HOMER-selected background.

### Alignment and preprocessing of functional genomic data for ABC score pipeline

Gestational week 18 (GW18) bulk ATAC-seq and H3K27ac ChIP-seq data from human fetal prefrontal cortex^64^ were aligned to hg19 using the standard Encode Consortium ATAC-seq and ChIP-seq pipelines respectively with default settings and pseudo replicate generation turned off (https://github.com/ENCODE-DCC). Trimmed, sorted, duplicate and chrM removed ATAC-seq and sorted, duplicate removed ChIP-seq bam files produced by the Encode pipeline were provided as input for calculating ABC scores.

ATAC-seq and H3K27ac CUT&RUN data from 7-8 week old NGN2-iPSC inducible excitatory neurons was obtained from Song 2019^29^. ATAC-seq and CUT&RUN reads were trimmed to 50 bp using TrimGalore^65^ with the command --hardtrim 5 50 before alignment. ATAC-seq reads were aligned to hg19 using the standard Encode Consortium ATAC-seq and ChIP-seq pipelines respectively with default settings and pseudo replicate generation turned off. Trimmed, sorted, duplicate and chrM removed ATAC-seq bam files from multiple biological replicates were combined into a single bam file using samtools merge v1.10^66^. Trimmed CUT&RUN reads were aligned to hg19 using Bowtie2 v2.3.5.1^67^ with the following settings --local --very-sensitive-local --no-mixed --no-discordant -I 10 -X 700 and output sam files were convert to bam format using samtools view ^66,67^. Duplicated reads were removed from the CUT&RUN bam file using Picard MarkDuplicates v2.26.0^68^ with the --REMOVE_DUPLICATES =true and -- ASSUME_SORTED=true options (http://broadinstitute.github.io/picard/). The final ATAC-seq and CUT&RUN bam files were provided as input for calculating ABC scores.

### Preprocessing of HiC and pcHiC data for ABC score pipeline

HiC contacts with 10 kb resolution from human GW17-18 fronto-parietal cortex was obtained in an hdf5 format separated by chromosome^69^(Supplementary Table 5). Hdf5 files were filtered for contacts with a score > 0 and converted into a bedpe format. Promoter capture HiC (pcHiC) contacts from 7-8 week old NGN2-iPSC inducible excitatory neurons were obtained in an ibed format from GSE113483^29^. The ibed file was converted to bedpe format and separated by chromosome. Bedpe files from GW17-18 cortex and iPSC derived excitatory neurons were provided as input for calculating ABC scores.

### Identification of candidate enhancer-gene pairs with ABC Score

The Activity-by-Contact (ABC) model identifies enhancer-gene relationships based on chromatin state and conformation^45^. Previously identified open chromatin regions from GW18 human prefrontal cortex^64^ and corresponding ATAC-seq and H3K27ac ChIP-seq bam files were provided as input for the ABC score pipeline MakeCandidateRegions.py script with the flags -- peakExtendFromSummit 250 --nStrongestPeaks 150000. Candidate enhancer regions identified were then provided to the run.neighborhoods.py script in addition to hg19 promoter merged transcript bounds. Finally, predict.py was used to identify final candidate enhancers using HiC data from human GW17-18 fronto-parietal cortex with the flags --hic_type bedpe --hic_resolution 10000 --scale_hic_using_powerlaw --threshold .02 --make_all_putative^69^. Candidate enhancer-gene pairs were also identified for 7-8 week old NGN2-iPSC inducible excitatory neurons using respective open chromatin regions^29^, ATAC-seq and H3K27ac ChIP-seq data. All other settings for the ABC score pipeline remained constant.

### Mouse enhancer transgenic assay

Transgenic E11.5 mouse embryos were generated as described previously^15^. Briefly, super-ovulating female FVB mice were mated with FVB males and fertilized embryos were collected from the oviducts. Regulatory elements sequences were synthesized by Twist Biosciences. Inserts generated in this way were cloned into the donor plasmid containing minimal Shh promoter, lacZ reporter gene and H11 locus homology arms (Addgene, 139098) using NEBuilder HiFi DNA Assembly Mix (NEB, E2621). The sequence identity of donor plasmids was verified using long-read sequencing (Primordium). Plasmids are available upon request. A mixture of Cas9 protein (Alt-R SpCas9 Nuclease V3, IDT, Cat#1081058, final concentration 20 ng/μL), hybridized sgRNA against H11 locus (Alt-R CRISPR-Cas9 tracrRNA, IDT, cat#1072532 and Alt-R CRISPR-Cas9 locus targeting crRNA, gctgatggaacaggtaacaa, total final concentration 50 ng/μL) and donor plasmid (12.5 ng/μL) was injected into the pronucleus of donor FVB embryos. The efficiency of targeting and the gRNA selection process is described in detail in Osterwalder 2022^15^. Embryos were cultured in M16 with amino acids at 37oC, 5% CO2 for 2 hours and implanted into pseudopregnant CD-1 mice. Embryos were collected at E11.5 for lacZ staining as described previously^15^. Briefly, embryos were dissected from the uterine horns, washed in cold PBS, fixed in 4% PFA for 30 min and washed three times in embryo wash buffer (2 mM MgCl2, 0.02% NP-40 and 0.01% deoxycholate in PBS at pH 7.3). They were subsequently stained overnight at room temperature in X-gal stain (4 mM potassium ferricyanide, 4 mM potassium ferrocyanide, 1 mg/mL X-gal and 20 mM Tris pH 7.5 in embryo wash buffer). PCR using genomic DNA extracted from embryonic sacs digested with DirectPCR Lysis Reagent (Viagen, 301-C) containing Proteinase K (final concentration 6 U/mL) was used to confirm integration at the H11 locus and test for presence of tandem insertions^15^. Only embryos with donor plasmid insertion at H11 were used. The stained transgenic embryos were washed three times in PBS and imaged from both sides using a Leica MZ16 microscope and Leica DFC420 digital camera.

## Contributions

M.Ko. conceptualized and designed the experiments and performed the analyses. D.L.C. conducted all MPRA-related experiments. M.Ko., D.L.C and N.A. wrote the manuscript. M.Ko., J.A.A., I.P-F., C.N., M.Ka., R.D.H, K.v.M., S.B., P.G., E.B. conducted transgenic mouse assay experiments. N.P. ran ABC analysis. I.G.S. ran TFBS analysis. S.S. provided computational resources for ABC analysis. N.A. and L.A.P. provided funding for the study. N.A. and L.A.P. supervised the study.

## Data Availability

All raw and processed data relating to the MPRA experiment can be found on the ENCODE portal under ENCSR865OZI, ENCSR517VUU, ENCSR548AQS and ENCSR257CZP. Data related to transgenic assays can be found at VISTA Enhancer Browser (enhancer.lbl.gov).

## Acknowledgments

This work was supported in part by the National Human Genome Research Institute (NHGRI) grant numbers UM1HG009408 (NA), UM1HG011966 (NA), R01HG003988 (L.A.P.), the National Institute of Mental Health (NIMH) grant numbers U01MH116438 (NA), R01MH109907 (NA) and National Institute of Child Health and Human Development (NICHD) grant number R01HD114353 (L.A.P). Part of the research was conducted at the E.O. Lawrence Berkeley National Laboratory and performed under U.S. Department of Energy Contract DE-AC02-05CH11231, University of California.

