## Supplementary Figures for "Massively parallel reporter assays and mouse transgenic assays provide complementary information about neuronal enhancer activity"

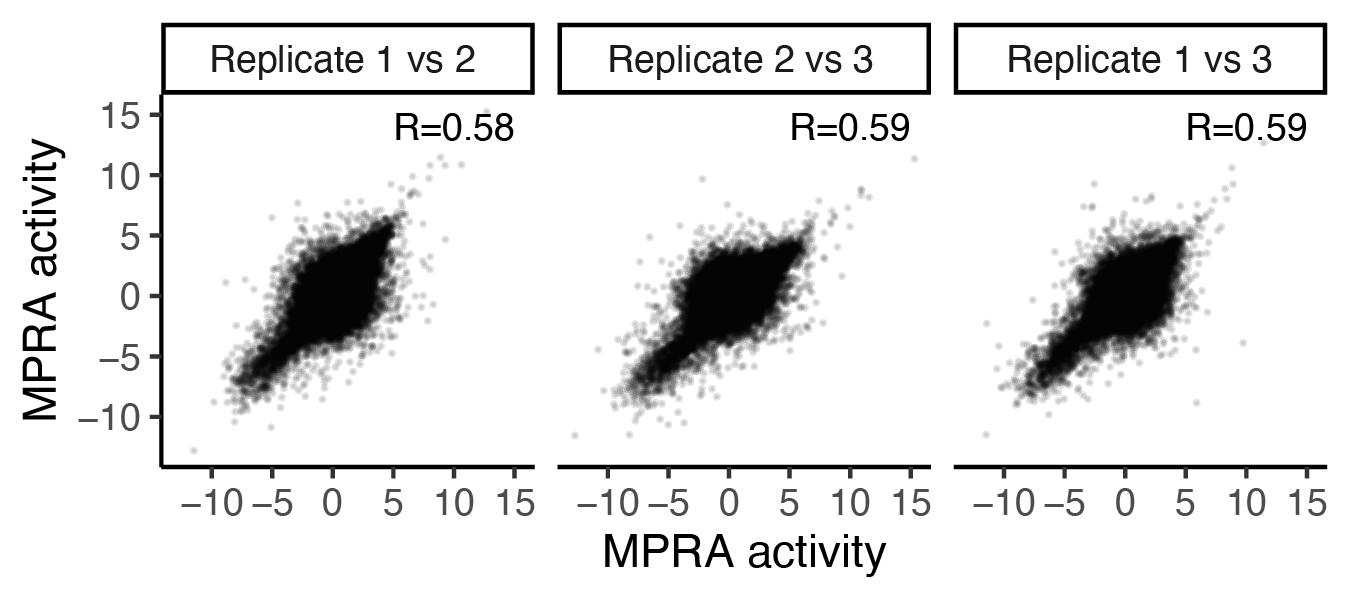

**Supplementary Figure 1**. Correlation of MPRA activity between biological replicates. R - Pearson correlation.

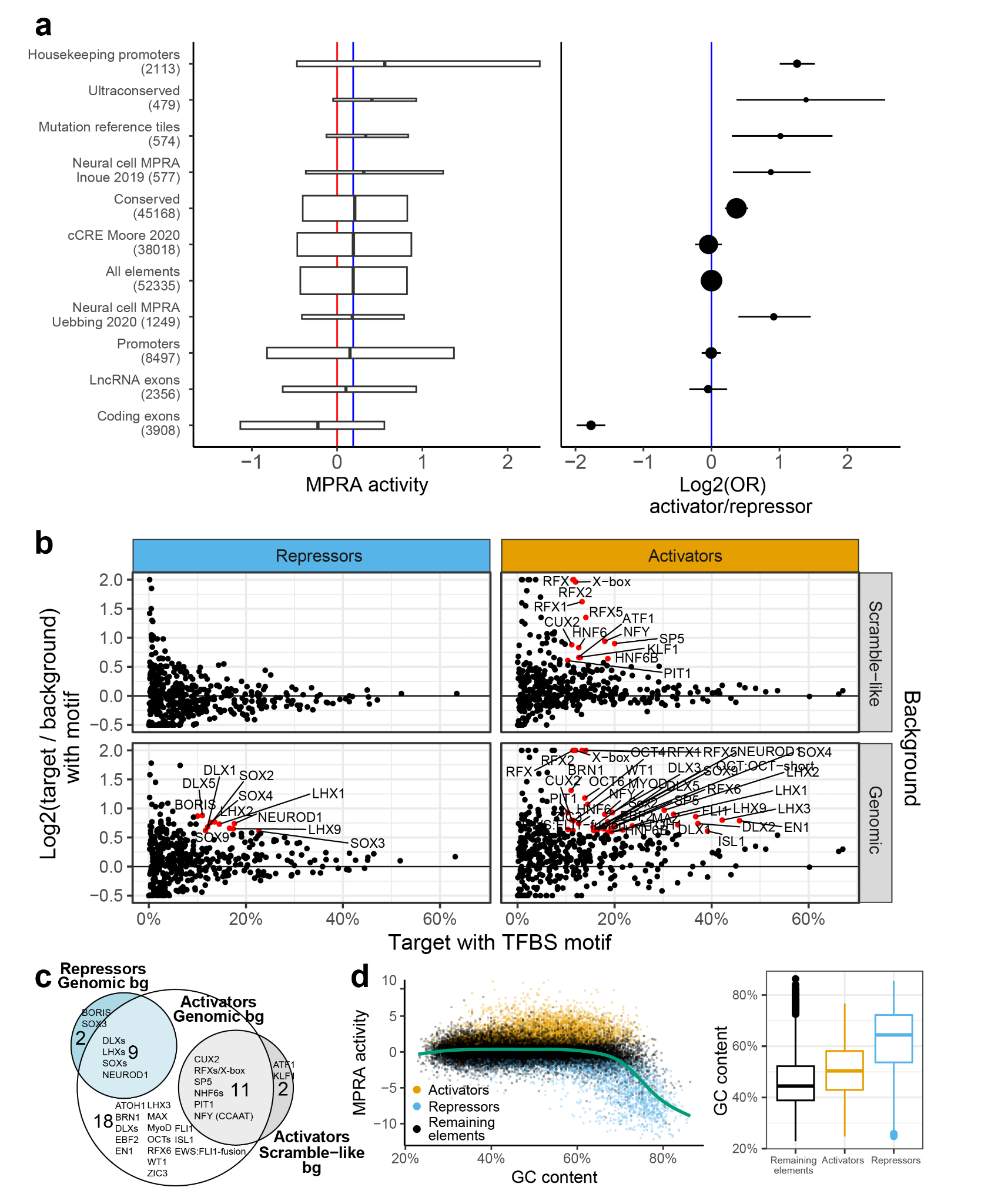

**Supplementary Figure 2**. **Neuronal WTC11 MPRA results validation. (a)** Activity (left) and enrichment of significant activator/repressor tiles (right) across functional categories. Red line = activity of scrambled negative controls (zero, by definition). Blue line = activity (0.19) or log-odds-ratio of activators to repressor tiles (zero, by definition) of all reference elements. All categories in left panel have significantly different activity than scrambled negatives at FDR-adjusted p-value < 0.05 (Mann-Whitney U test). Bars in log-odds plot in the right panel are 95% confidence intervals - all categories with interval not overlapping 0 are significant at FDR-adjusted p-value<0.05 (Fisher test). See Supplementary Table 1 for source data. Left panel is the same as Figure 2a. **(b)** TFBS enrichment in enhancer activator (N = 3,054) or repressor (N = 1,894) tiles compared to enhancer elements with scramble negative levels of activity (N = 15,503, "scramble-like") or genomic background elements (N = 50,000; "genomic"). Log2-fold change was curbed at -0.5 and 2. Only TFBSs present in more than 10% target, with 50% increase in presence from background to target set (corresponding to log2(1.5) = 0.58 cutoff) and FDR<1% are labeled. Top-right panel is the same as Figure 2b. **(c)** Overlap between TFBS enriched in different analyses from previous panel. TFs with similar names collapsed (e.g. "RFXs"). **(d)** Relationship between MPRA activity and GC content. Left: scatterplot. Green line is a smooth mean generated by a general additive model with REML parameter selection. Right: boxplot of GC content binned by tile category (activators N = 4,762, repressors N = 2,957, remaining elements N = 44,616). Hinges of boxplots span interquartile range (IQR), line in the middle is median, thickness (height) is proportional to number of overlapping tiles. Where used, whiskers extend from the hinge to the largest value no further than 1.5 * IQR from the hinge.

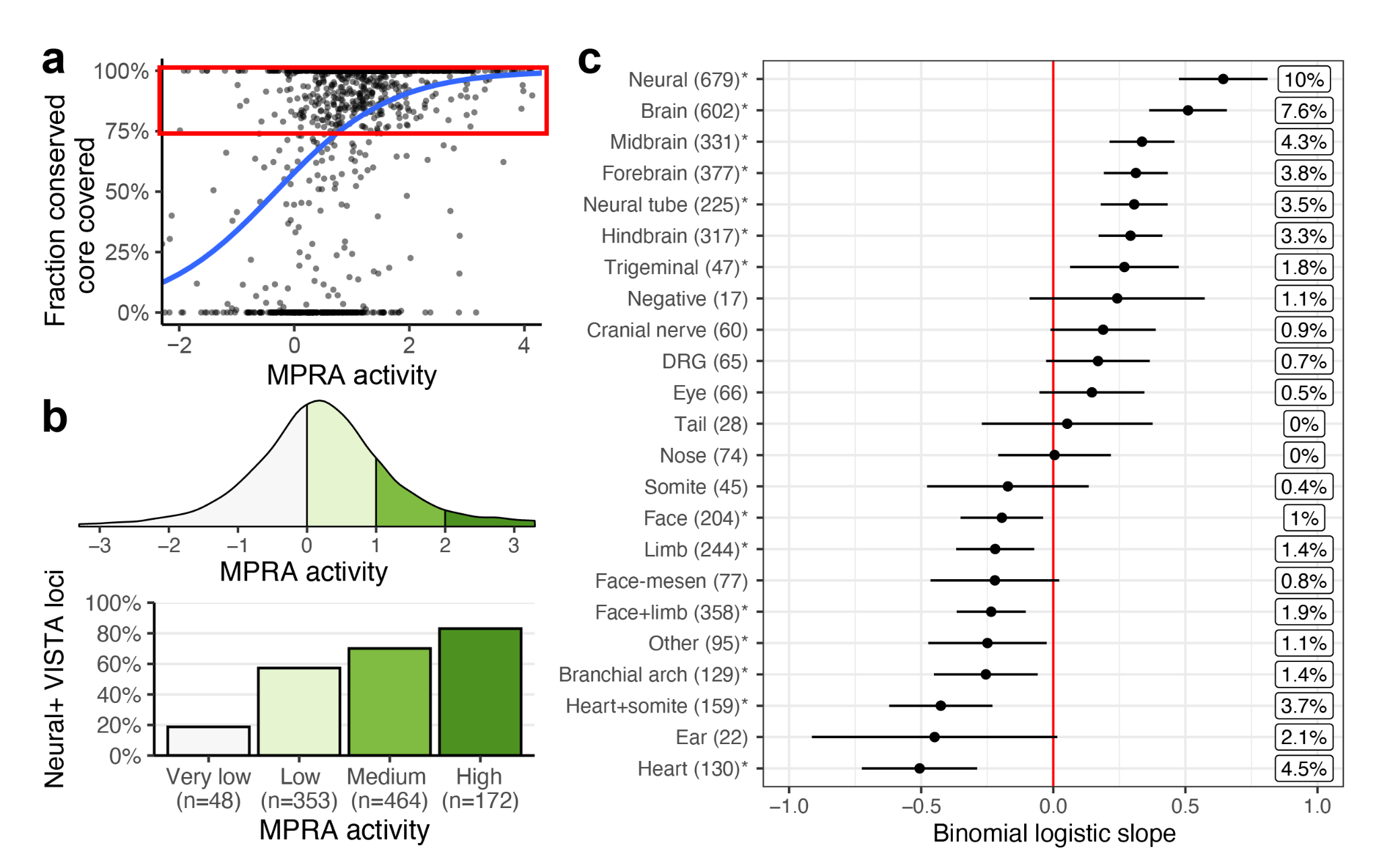
**Supplementary Figure 3**. **Predicting transgenic assay activity using a MPRA-based, coverage-marginalized model. (a)** Relationship between fraction of conserved core covered and MPRA activity. The blue line is the binomial-link GLM regression on this variable. Instead of including this variable as covariate in GLM, VISTA elements with coverage lower than 75% were removed prior to modeling. **(b)** Alternative visualization of relationship between MPRA activity and transgenic assay activity. Top: MPRA activity bins. Bottom: fraction of neural-positive VISTA loci by MPRA activity bin. Numbers below bars are counts of VISTA elements. Only "well-covered" elements included, as defined above (N = 1,037). **(c)** Results of the GLM predicting binomial transgenic assay activity of well-covered VISTA elements from MPRA activity. Asterisks indicate nominal p-value < 0.05. Boxed percentages to the right are Nagelkerke R^2^ measures. DRG = dorsal root ganglia. Face-mesen = facial mesenchyme. Cranial nerves category does not include the trigeminal nerve, as per VISTA Browser.

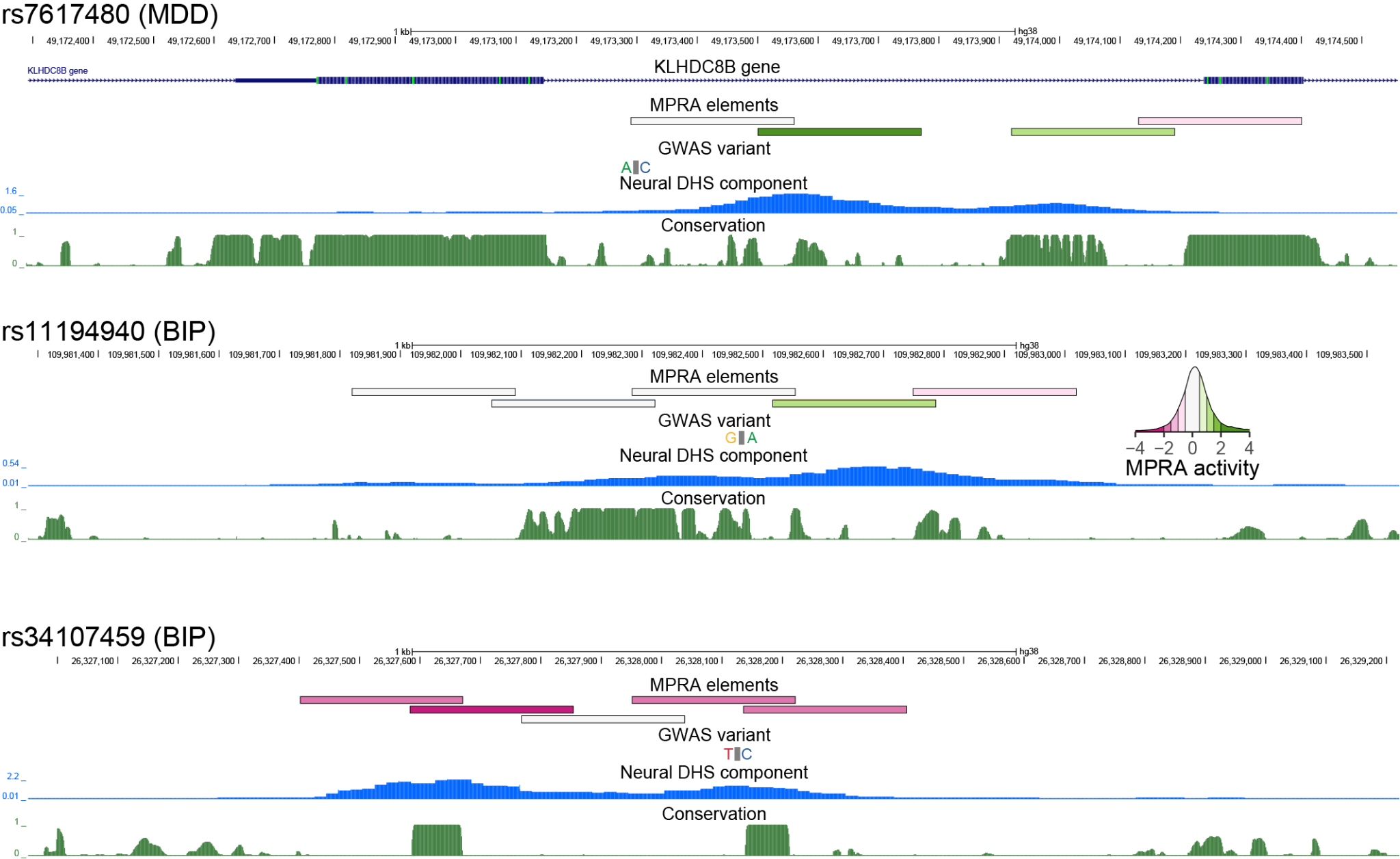

**Supplementary Figure 4**. **Genomic tracks of the three nominally significant GWAS variants.** Conservation is PhastCons UCSC tracks for 30 mammals (27 primates) dataset. MPRA elements colored by MPRA activity, see inset. Neural DHS component from Meuleman 2020^32^. MDD = major depressive disorder. BIP = bipolar disorder.

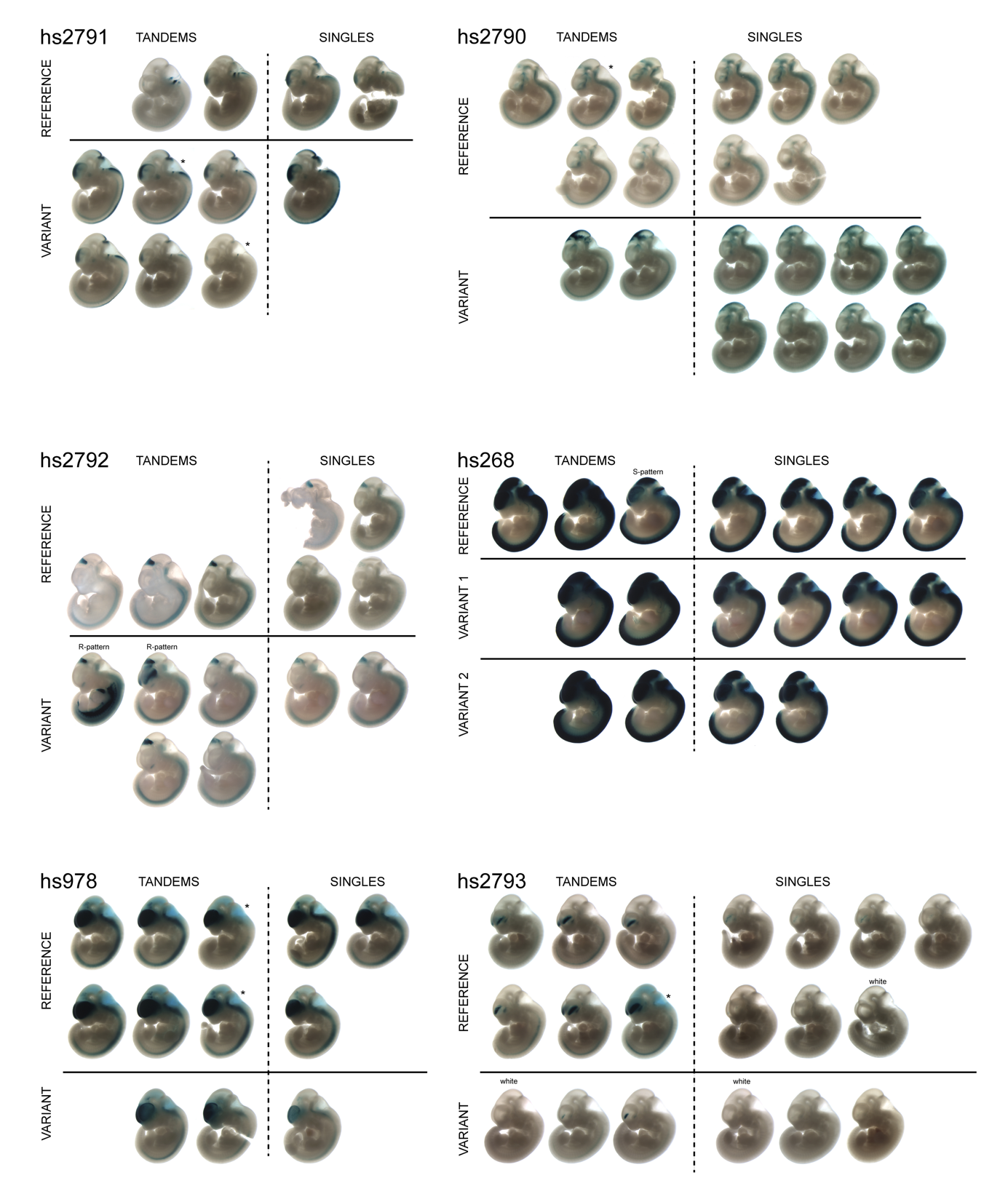

**Supplementary Figure 5. Results of transgenic mouse assay.** Tandems = embryos that were genotyped as positive for reporter integration at the safe harbor locus and presence of the plasmid backbone indicating higher transgene copy number with strong, reproducible pattern. Singles = embryos that were genotyped positive for reporter integration at the safe harbor locus and negative for plasmid backbone, indicating lower transgene copy number with weaker, but reproducible pattern. Asterisks - embryos with uncertain genotype. R-pattern - embryos with deviant pattern indicative of random (R) genomic insertion. S-pattern - tandem embryos with expression pattern resembling that of single-genotyped embryos. Embryos without any staining are marked as "white".

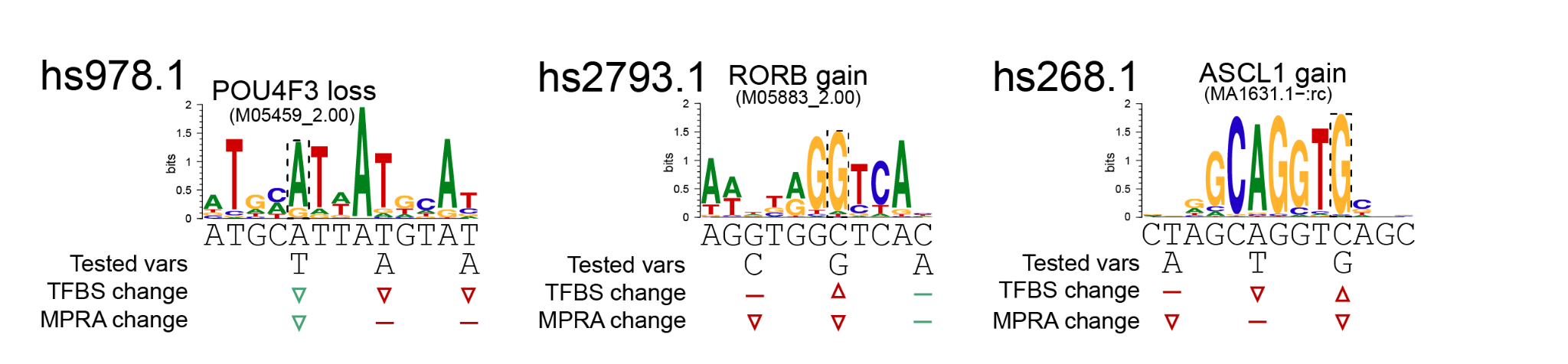

**Supplementary Figure 6.** Prediction of TFBS likely affected by the variants, but inconsistent with flanking variant effects.
